# Towards Semantic fMRI Neurofeedback: Navigating among Mental States using Real-time Representational Similarity Analysis

**DOI:** 10.1101/2020.11.09.374397

**Authors:** Andrea G. Russo, Michael Lührs, Francesco Di Salle, Fabrizio Esposito, Rainer Goebel

**Affiliations:** Department of Political and Communication Sciences, University of Salerno, Fisciano (Salerno), Italy; Department of Medicine, Surgery and Dentistry, Scuola Medica Salernitana, University of Salerno, Baronissi (Salerno), Italy; Department of Cognitive Neuroscience, University of Maastricht, Maastricht, The Netherlands; Brain Innovation B.V., Maastricht, The Netherlands

**Keywords:** neurofeedback, representational similarity, real-time fMRI, semantic representation

## Abstract

**Objective:** Real-time functional magnetic resonance imaging neurofeedback (rt-fMRI-NF) is a non-invasive MRI procedure allowing examined participants to learn to self-regulate brain activity by performing mental tasks. A novel two-step rt-fMRI-NF procedure is proposed whereby the feedback display is updated in real-time based on high level (semantic) representations of experimental stimuli via real-time representational similarity analysis of multi-voxel patterns of brain activity.

**Approach:** In a localizer session, the stimuli become associated with anchored points on a two-dimensional representational space where distances approximate between-pattern (dis)similarities. In the NF session, participants modulate their brain response, displayed as a movable point, to engage in a specific neural representation. The developed method pipeline is verified in a proof-of-concept rt-fMRI-NF study at 7 Tesla using imagery of concrete objects. The dependence on noise is more systematically assessed on artificial fMRI data with similar (simulated) spatio-temporal structure and variable (injected) signal and noise. A series of brain activity patterns from the ventral visual cortex is evaluated via on-line and off-line analyses and the performances of the method are reported under different noise conditions.

**Main results:** The participant in the proof-of-concept study exhibited robust activation patterns in the localizer session and managed to control the neural representation of a stimulus towards the selected target, in the NF session. The offline analyses validated the rt-fMRI-NF results, showing that the rapid convergence to the target representation is noise-dependent.

**Significance:** Our proof-of-concept study demonstrates the potential of semantic NF designs where the participant navigates among different mental states. Compared to traditional NF designs (e.g. using a thermometer display to set the level of the neural signal), the proposed approach provides content-specific feedback to the participant and extra degrees of freedom to the experimenter enabling real-time control of the neural activity towards a target brain state without suggesting a specific mental strategy to the subject.

## 1. INTRODUCTION

Real-time functional magnetic resonance imaging neurofeedback (rt-fMRI-NF) is a psychophysiological approach in which the on-line measured blood oxygen level dependent (BOLD) signal is provided to the subject as visual [1,2], auditory [3], haptic [4] or electrical [5] feedback to allow the self-regulation of his/her own neural activity towards target levels [6]. The increasing performance in this task has been previously associated with measurable improvements in specific neurological functions and/or positive changes in behaviours [7], thereby rt-fMRI-NF has been successfully applied in a great variety of domains such as motor function [8,9], emotion regulation [10,11], prosody [12] and visual task performance [13]. Furthermore, rt-fMRI-NF has found applications in the treatment of different neuropsychiatric disorders [14–18].

The flexibility of fMRI as a functional neuroimaging tool and its successful integration with advanced computational and real-time analysis methodologies has contributed to the development of various experimental rt-fMRI-NF frameworks [19]. However, there is an ongoing debate on whether or not participants in rt-fMRI-NF sessions need to be provided with an explicit strategy to regulate their own brain activity. Indeed, both approaches have their strengths and weaknesses and, in most cases, the choice of one over the other depends on the research question [19]. Thus, there are studies that provide the subjects with a clear strategy [8,20,21] as well as studies in which the subject is left free to choose the task [3,7,13,22]. Furthermore, there are various choices for the source of the NF signal and how it can be calculated. For example, some studies have used the variation in the average fMRI signal from a single region of interest (ROI) compared to a baseline [17,23,24], whereas others rely on (a measure of) the functional connectivity within a network of two or more ROIs [25] to target more complex cognitive functions [7,26,27]. Estimating the likelihood of a decoded neural activity pattern starting from a previously recorded brain pattern, is also possible [7,13,28]. Finally, there is a wide spectrum of viable options also concerning the feedback modalities (e.g. visual, auditory, haptic and electrical) and how to present the NF signal to the subject. For instance, visual feedback can be presented using simple shapes (e.g. a vertical bar or a circle) [29] as well as more immersive virtual reality interfaces [2].

However, although different NF approaches have been successfully applied in several domains [7,11,14,15,19], the possibility to provide a more semantically-driven feedback, that, by some means, encodes the current mental state of the subject from high-level brain representations, as well as its differences from previous and other mental states, could be critically useful in some challenging scenarios such as emotion regulation [11]. Along these lines, a semantic representation of the stimulus estimated with the application of a multidimensional approach to the current participant’s brain activity may provide the extra degrees of freedom to self-modulate a mental state.

The aim of this work is to propose a novel rt-fMRI-NF paradigm based on a real-time incremental version of representational similarity analysis (RSA) [30], referred to as real-time RSA (rt-RSA). RSA allows an abstract representation of brain activity patterns, in terms of concepts and associated mental states modulated by a given task. Thus, the core of this approach is the creation of a visual feedback display that is related to the similarity space of a participant’s current brain activity pattern in relation to patterns from a set of related mental tasks measured during localizer runs. The patterns are extracted from chosen brain regions that represent semantic information related to the goals of an NF study (e.g. related to visual categories or different emotions). In its real-time implementation, the semantic feedback display represents the subject’s current neural activity as a movable point in a plane where a set of candidate target neural states are displayed as fixed points. The feasibility and validity of the proposed approach have been tested on real and artificial data.

## 2. MATERIALS AND METHODS

### 2.1 Representational Similarity Analysis

Representational theory [31] can help neuroscientists in interpreting the spatial distribution of brain activity across several neuro-physical units (e.g. neurons or voxels). It is assumed that, in a given time window of observation, the spatial distribution or pattern encodes the neural representation of contents of, e.g., an image, a sound, or a motor action. On these premises, the combination of different patterns of neuronal activity defines a multi-dimensional space where the dimensions are the neuro-physical units and a single point in this space is a pattern [32]. The geometrical properties of this space can be characterized and analysed using RSA [32]. For example, considering a set of experimental conditions and their corresponding brain-activity patterns from an ROI, it is possible to estimate their mutual (dis)similarities and to encode these in a representational dissimilarity matrix (RDM) [30,32,33]. The RDM operationally defines the RS of the selected region and can be used to compare different representations (from different brain regions or conditions) via a signed-rank test [30,33,34]. The most common measure to compute the dissimilarities between two brain patterns is the correlation distance (i.e. one minus the linear correlation between spatial patterns). Statistically, this measure normalizes for both the mean and the standard deviation of the spatially variable activity. Geometrically, it is related to the angle between two high dimensional vectors (i.e. the brain patterns) and ranges from a minimum of 0 to a maximum of 2: if the two patterns are highly correlated, their dissimilarity goes to 0, whereas if they are anti-correlated, this value goes to 2. Other possible measures are the Euclidean distance, the Mahalanobis distance, the absolute activation differences and a computational measure that is based on the pairwise classification accuracies of discriminative models (e.g. via linear discriminant analysis) [30,33,35].

Using a dimensionality reduction approach like multidimensional scaling (MDS) [36–38], the RDM can be visualized in a two or three-dimensional scatter plot where the mutual distances among points reflect the dissimilarities among the response patterns [30,33].

RSA has gathered important insights in various domains, including vision [39,40], audition [41], categorical perception [42,43], memory [44] and motor control [45]. For a more exhaustive review please see [32].

### 2.2 Real-time RSA

The rt-RSA approach integrates RSA into an NF paradigm enabling the comparison of multiple neural patterns (representing a given set of experimental conditions) between each other and intuitively summarizing their differences [32] in real-time. In its simplest version, for a given ROI, a visual feedback is provided to the subject consisting of a constellation of points anchored on a plane, called “representational space” (RS), corresponding to the target neural representations of a set of base stimuli, and a moving point corresponding to a variable neural representation. The coordinates of these points on the plane are estimated (and eventually those of the moving point updated) from multi-voxel patterns of regional BOLD activation which have been previously shown to identify (or encode the dynamic variability of) mental (e.g. cognitive or emotional) states. Thereby, following the initial estimation of the RS from an fMRI localizer session, the participant’s ROI activity pattern will be estimated in real-time and provided as a novel movable point in this space. The aim is to let the participant *move* this point towards a selected target point in the RS by learning to self-modulate his/her own multi-voxel activation pattern in such a way to engage in a specific mental state. To this purpose, the participants are provided intermittently with a visual feedback showing the position of their current brain state within the RS with respect to the base stimuli.

### 2.3 Mathematical description of the method

The possibility to project a newly calculated brain activity pattern on the existing (previously estimated) RS is based on the application of a linear solution to the distance-based triangulation problem, introduced in the field of MDS by de Silva and Tenenbaum (2004). The solution was part of a different multidimensional scaling approach originally proposed to overcome the limitation of classical multidimensional scaling (cMDS) [37,46], also known as Principal Coordinate Analysis (PCoA), when the number of entries of the input matrix is very large compared to the intrinsic dimensionality of the data (for a complete description of the method, please see [47]). The cMDS belongs to the family of MDS methods, as it projects elements from a high-dimensional space (e.g. a brain pattern) to a lower-dimensional space while preserving the geometry as faithfully as possible [47].

The solution for the distance-based triangulation problem provides a convenient mathematical framework to project a new data point (i.e. a new brain pattern) onto an existing RS by simply applying a previously estimated linear transformation. Specifically, given a set of base stimuli and their evoked activity patterns in an ROI, an RDM is initially calculated based on their pairwise dissimilarities. In our case, the metric chosen for this calculation is the correlation distance. Then, the RS corresponding to the estimated RDM is obtained by applying the cMDS to the RDM itself. Finally, using a fixed linear transformation function, the coordinate vector *<u>x</u>* representing the position of the pattern of a new stimulus in the estimated RS is obtained according to the following formula:

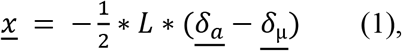

where

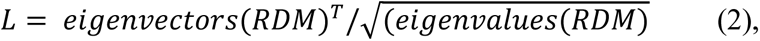

*<u>δ_a_</u>* is the squared vector of the dissimilarities between the new stimulus and the base stimuli and <u>*δ*_μ_</u> is the sum of the squared columns of the RDM divided by its number of entries (i.e. the number of base stimuli).

### 2.4 General experimental framework

An rt-fMRI-NF experiment with the use of rt-RSA requires at least two separate sessions: a localizer session and an NF session. To increase the statistical power of the estimated brain activity patterns, the localizer session may include multiple runs where each run can include several trials of one base stimulus (one run per base stimulus) or interleaved trials of different base stimuli. In each localizer session, a series of N trials (corresponding to different repetitions of each base stimulus) are delivered to the subject who performs an active (e.g. button pressing) or passive (e.g. image viewing) task. The ROI is ultimately selected at the end of the localizer eventually using a combination of functional and anatomical criteria based on the measured brain activation and a priori regional hypotheses, if available [35].

A single activation pattern (for a given stimulus) can be obtained from the ROI map of regression coefficients (as effect size estimates, see, e.g., [35]) or statistical parameters (e.g. t scores, signal-to-noise estimates, see, e.g., [35]) via the general linear model (GLM) analysis of the fMRI responses at each voxel. The GLM estimation can be either performed online, thereby the signal (and noise) incremental estimates are recursively updated over successive trials, or offline, by eventually concatenating multiple runs using the most complete basis set of stimulus and confound predictors, to maximize the accuracy and power of all base stimulus patterns (to be used for the base RS).

During the NF session, the multi-voxel pattern of the selected ROI is updated via online GLM and its position within the previously estimated RS is periodically displayed as a visual stimulus to the subject. The participants are thus periodically informed about the position of their current mental representation (as obtained from their current brain activity) with respect to the mental representations associated with all base stimuli (as obtained from the brain activity measured during the localizer runs). In this way, the subjects are stimulated to self-modulate their own brain activity to change its position within the RS towards the target position. Operationally, at the end of each task block, the participant’s multi-voxel pattern is extracted from the predefined ROI and the dissimilarities of this pattern vs. the base stimulus patterns are calculated and given as input to (1) to update the corresponding coordinates in the RS. Moreover, while collecting the series of new patterns (i.e. mental states) over successive NF trials, the trajectory of the current mental state is also displayed to the subject, thereby the history of the modulation is kept visible to the subject to possibly incentive (or disincentive) an undertaken strategy towards reaching the target state.

A graphical description of the procedure is provided in figure 1.

**Figure 1.**
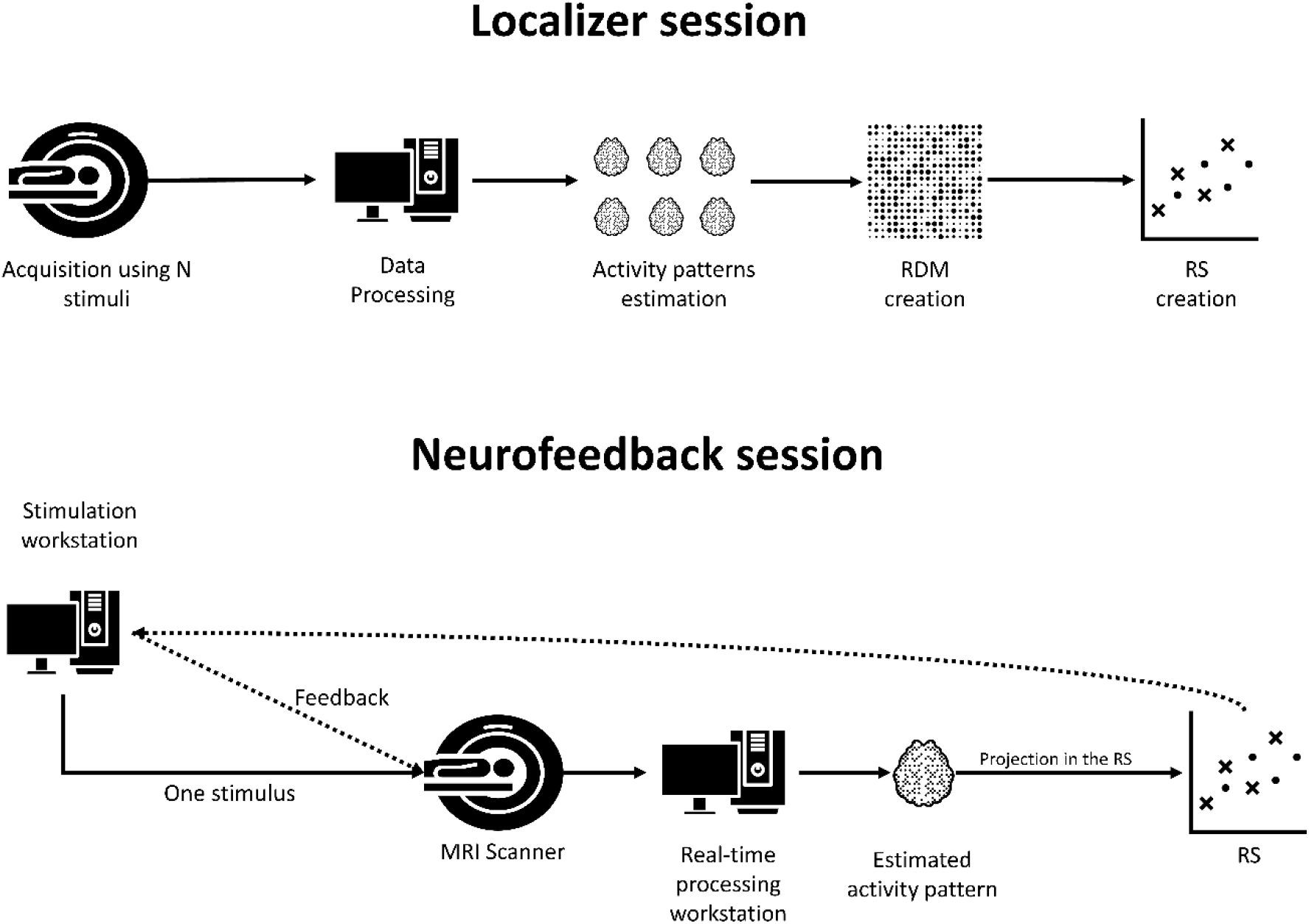
General workflow of an fMRI-NF experiment with the use of the rt-RSA. In the localizer session the subjects perform a task (e.g. passive viewing) and are exposed to a set of experimental conditions. The brain-activity related to each experimental condition is extracted from a region-of-interest (ROI) and used to estimate a representational dissimilarity analysis (RDM). Then, the latter is projected on a plane that is the representational space (RS) of the ROI. In the neurofeedback session, the subjects perform the same task but are provided with one stimulus. The fMRI data are processed in real-time and the brain activity of the ROI (used in the localizer session) are periodically extracted and projected in the RS. The position of this projection in the RS is provided intermittently to the subject as visual feedback.

### 2.5 Method implementation

The proposed method was implemented in Python (Python Software Foundation. Python Language Reference, version 3.7. Available at http://www.python.org)(van Rossum, 1995). The fMRI data were processed online and offline using Turbo-BrainVoyager version 4.0 (TBV) (Brain Innovation B.V., Maastricht, The Netherlands) and imported in Python using the TBV Network Access Plugin (<u>Network Access Plugin</u>) and the corresponding interface (<u>GitHub - expyriment/expyriment-stash</u>) as part of the open-source library Expyriment [48]. All the scripts and the data used in this work are available at https://github.com/andreagrusso/rtRSA.

### 2.6 fMRI Neurofeedback experiment

#### 2.6.1 Participant

Functional and anatomical data of one healthy left-handed volunteer (male, age 27), with normal vision and without known neurological or psychiatric disorders, were obtained using a 7T scanner (Siemens Healthcare, Germany). Informed written consent was obtained before the study and the experimental procedure was approved by the local Ethics Committee of the Faculty of Psychology and Neuroscience at Maastricht University. The experimental procedure conformed to the principles embodied in the Declaration of Helsinki.

#### 2.6.2 Experimental design

The complete scanning session was divided into two main parts: an offline training session (outside the MR scanner) and a scanning session including the localizer and the NF experiment. In both experiments (localizer and NF) the participant performed an imagery task upon the delivery of an auditory cue.

In the training session, the subject was asked to familiarize with the stimuli by visually inspecting and memorizing the images of selected objects as well as listening to their accompanying cues. Two animate (cat, dog) and two inanimate (chair, hammer) objects were chosen as base stimuli. The training continued inside the MR scanner during the acquisition of the anatomical data.

The localizer experiment was composed of four consecutive functional runs, one for each object. A single run was experimentally designed as a series of ten tasks and eleven rest blocks of 20 seconds in which the participant alternately imagined the object corresponding to the delivered auditory cue (e.g. “cat” or “chair”). At the end of each task-block, the word “stop” was delivered to the participant to announce the beginning of the resting period. The subject was requested to focus the gaze on a white fixation cross centred on a dark grey screen during the whole acquisition. The four images were selected from a database of naturalistic objects [49] while the corresponding auditory cues were generated using https://soundoftext.com/ that creates audio files from text using the text to speech engine of Google Translate. Both the auditory cues and the visual feedback were delivered to the subject using a custom-made Python script using PsychoPy3 module [50].

The NF run was designed according to a similar experimental paradigm. Specifically, the subject was requested to imagine for a period of 20 seconds the object upon delivery of the corresponding auditory cue. At the end of the task-block, the subject heard the word “stop” that indicated the beginning of a 20 second period of rest. However, differently from the training and localizer runs, the rest-period was followed by an extra-period of 5 seconds during which the visual feedback was displayed. This consisted in the RS display where the positions of the base stimuli (i.e. the activity patterns acquired in the localizer session) were displayed as yellow points with a label tag indicating the object name. The position of the *current* activity pattern, as estimated from the last time window of measurement (40 s) ending right before the feedback-block, was displayed as a red star. In addition, the positions of the *current* pattern from all previous task-blocks were also displayed as light-grey stars and linked to one another with a green line. Thereby, the participant was provided, not only with the position in the RS of the *current* brain pattern but also with the trajectory performed across the previous task-blocks. The experimental procedure is summarized in figure 2.

**Figure 2.**
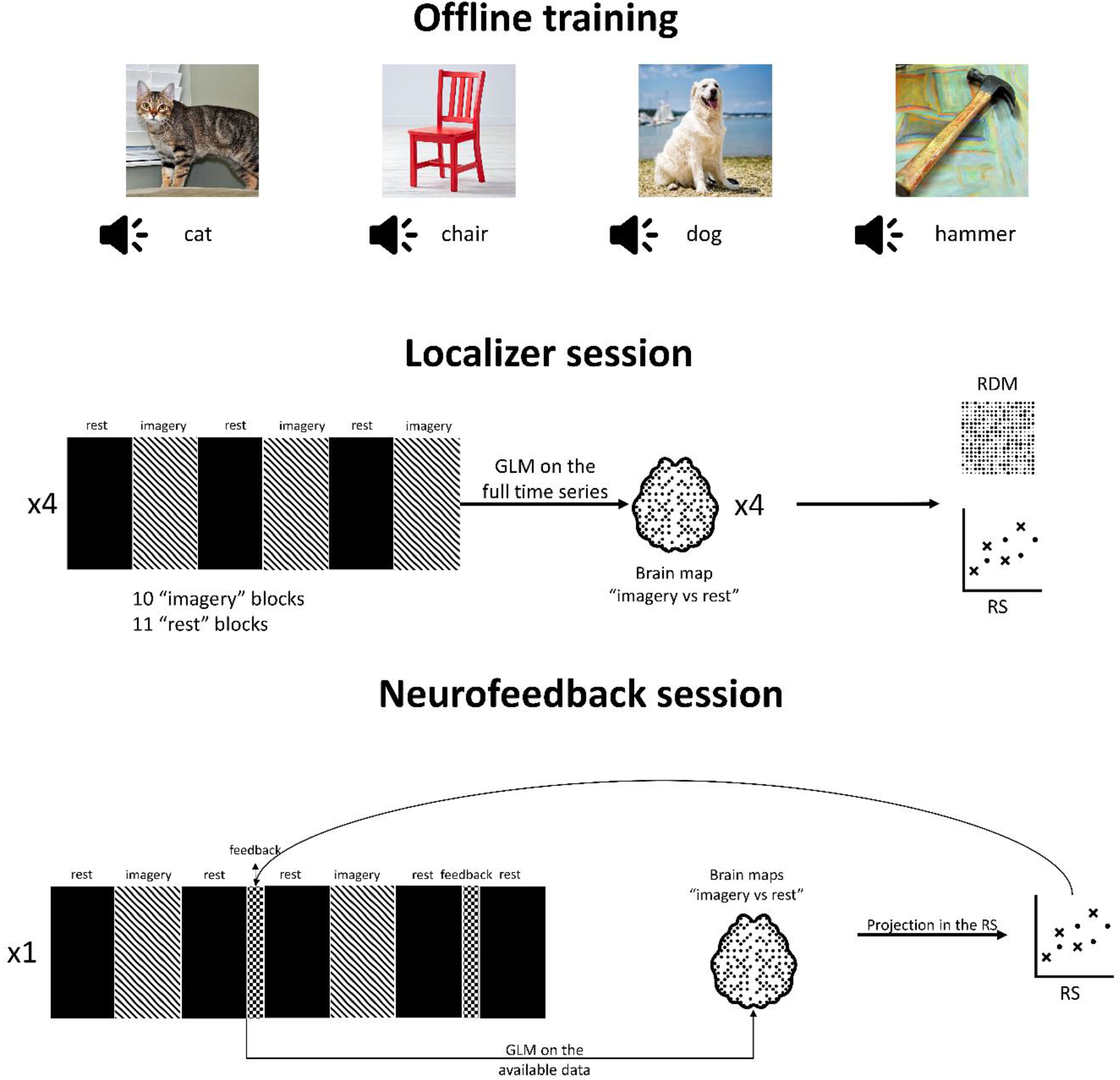
Workflow of the fMRI-NF experiment performed. In the offline training session, the subject was requested to memorize four images and their corresponding auditory cues. In the localizer session, the subject underwent four block-design fMRI acquisitions, one for each stimulus, in which he was requested to imagine as clear as possible the image corresponding to the provided auditory cue. The imagery blocks were interleaved by periods of rest. At the end of each functional acquisition the statistical values relative to the contrast “imagery vs rest” was extracted for all the voxels of a defined ROI. The dissimilarities among these activation patterns were estimated and first, encoded in a RDM and projected onto a plane to create the RS of the ROI. In the neurofeedback session the subject was provided with only one auditory cue and requested to imagine the corresponding image. The imagery blocks were interleaved by periods of rest and small period of feedback. At the beginning of the feedback period the activity pattern of the defined ROI relative to the contrast “imagery vs. rest” is extracted and projected in the RS space. The RS with the new point is given as feedback to the subject.

#### 2.6.3 Data acquisition

FMRI data were obtained using a 7T whole-body scanner (Magnetom; Siemens AG, Erlangen, Germany). The participant was placed comfortably in the MRI scanner with foam padding next to the head to minimize spontaneous or task-related motion.

For the functional acquisition, multi-band [51–53] repeated gradient-echo echo-planar imaging (EPI) sequences were used. Except for the number of time points (localizer session: 430 volumes; NF session: 680 volumes), identical scanning parameters were used for all functional measurements (repetition time (TR) = 1000 ms, echo time (TE) = 21 ms, number of axial slices = 60, matrix = 112 × 112, field of view (FOV) = 224 mm, thickness = 2 mm, interslice gap = 0 mm, multi-band factor = 4). In the NF session, functional images were reconstructed and exported using a direct TCP/IP connection from the image reconstruction computer to the real-time analysis computer and stored on the hard drive. The real-time data analysis software (TBV) running on the real-time analysis computer was able to read and process the exported images in real-time.

For the acquisition of the structural data, a high-resolution T1-weighted anatomical scan was acquired using a three-dimensional (3D) magnetization prepared rapid-acquisition gradient-echo (MP2RAGE) sequence (192 slices, 0.9 mm iso. voxels, no gap, TR = 4500 ms, TE = 2.39 ms, TI1 = 900ms, TI2 = 2750, FA = 5, FOV = 230 × 230mm2, matrix size = 256 × 256, total scan time = 8 min and 34 s.

### 2.7 Online data processing

The functional runs of the localizer were processed with TBV to optimally define the target ROI to be used for the extraction of the brain activity patterns elicited by the base stimuli and for the subsequent NF runs. During the localizer session, the online data processing allowed the real-time monitoring of the subject’s head motion and the general quality of the data. More, specifically, motion correction with sinc interpolation, linear trend removal, temporal high-pass filtering and spatial smoothing with an isotropic 4-mm full-width at half-maximum (FWHM) Gaussian kernel were applied online to the functional time-series. The online voxel-wise GLM was computed incrementally with one predictor time-course for the imagery task (derived from an incremental boxcar function convolved with a standard hemodynamic response function [54]) and six motion parameters (incrementally derived from the motion correction procedure) as confound predictors. The online (incremental) GLM was fitted at each voxel using a recursive least squares estimation of the regression coefficients. At the end of the session, pre-processed functional data were reloaded in TBV and the offline GLM analysis was also applied to the complete pre-processed fMRI time series. In both cases, the t-contrast “imagery vs. rest” was calculated.

Starting from offline GLM results, an ROI was defined in the inferior temporal cortex (ITC) as this region is well known to encode high-level (semantic) representations of natural objects at the interface between vision and semantics [42,55,56]. For this study, the ROI definition was performed using a combined anatomical and functional approach. Namely, a whole-brain probabilistic functional map in MNI space was generated from the Neurosynth database [57] using the keyword “object”. The statistical threshold was set to q=0.1 using the False Discovery Rate [58]. This map was imported in BrainVoyager 21.4 (Brain Innovation, Maastricht, The Netherlands) and an ROI was initially defined by selecting a cluster of activation encompassing the left ITC. Before the NF run, the ROI definition was further adapted to the estimated brain activity from the offline GLM of localizer runs. Namely, the extracted ROI was imported in TBV where it was transformed to the native space of EPI images and used as a guide to manually define a functional ROI in the ITC from the offline GLM contrast “imagery vs. rest” (main effects of all stimuli, p<0.001). At the end of each functional run, for each voxel of the selected ROI, the t-values relative to the contrast “imagery vs. rest” were extracted and stored to local disk. Upon completion of all the functional runs of the localizer sessions, the stored data were assembled in a matrix, whose dimensions were the number of stimuli and the number of voxels of the ROI, and which was used to estimate the RDM and the corresponding RS (with its transformation) to be used in the NF session.

The data of the NF were processed in real-time with TBV with the same preprocessing steps used in the localizer session. The ROI activity pattern related to the contrast “imagery vs. rest” was extracted for the incremental time window ending one time point before the beginning of the feedback block, thereby encompassing the whole time series up to this point in time. The incremental GLM included the estimated motion predictors as confounds. To generate the feedback stimulus at the beginning of the feedback block, the extracted t-values from the ITC ROI were used to estimate the position of the current brain state in the RS estimated from the localizer data.

### 2.8 Offline data analysis

The full time-series data acquired during both the localizer and the NF session were reloaded in TBV and used for further analyses. Namely, the data from the localizer session were used to assess the computational and statistical performances of the rt-RSA approach on both real and artificial fMRI time-series, whereas the data from the NF session were used to analyse the experimental performance of the subject. For the validation, using the same GLM predictor as ideal response time-course, an artificial dataset with simulated brain activity was created to analyse the performances of the rt-RSA approach under different signal-to-noise conditions. In this way, the same incremental GLM analysis, as implemented in TBV, was used for the simulations.

#### 2.8.1 Stability of the rt-RSA

The data of the functional runs of the localizer session were reloaded in TBV. Starting from this data set, four NF experiments were simulated to investigate the stability of the rt-RSA approach. Namely, the t-maps from the incremental GLM were first extracted at the end of each task-block and used to simulate the dynamically updated brain pattern as the input of a virtual NF display. This pattern was projected as an additional point onto the RS estimated from the complete time-series of all functional runs. The distances, and the trajectory, from the input to the target are estimated at each block. As both online (incremental) and offline GLM analyses are conducted on the same localizer data, this scenario simulates the ideal successful NF outcome whereby the input NF pattern (from the online GLM) overlaps perfectly to the pattern associated with one of the base stimuli. Thereby, the stability of the rt-RSA can be evaluated.

#### 2.8.2 Performance evaluation

To evaluate the performance of the subject, both the localizer and the NF data were analysed. Using the localizer data, an RDM was calculated at the end of each task-block via the incremental GLM. The last RDM (i.e. the one estimated from the full time-series) was chosen as reference RDM and the monotonic correlation between the vectorized upper triangular part of the RDM at each block and the vectorized upper triangular part of the reference RDM was statistically evaluated with a signed-rank test [30,33,34]. The analysis of the RDM correlation series allowed us to have a possible estimate of the minimum number of blocks needed for the subject to incrementally generate an RDM which is not significantly different from the reference RDM. Moreover, it also allows the evaluation of the ability of the subject to maintain this similarity, consistently over time, across the subsequent blocks. The idea behind this analysis is that with an increase in the number of time points and, as a consequence, in the number of task blocks, it is possible to understand if the brain activity has become more stable and if the participant has been able to modulate consistently his brain activity. Therefore, this analysis could help us to evaluate prospectively, before the NF session, the quality of the localizer data as well as the stability of the base RS.

Using the data of the NF session, the performance of the participant was evaluated by measuring the distances in the RS of the projections of *current* brain patterns to the corresponding target. This analysis allowed us to have a measure of the participant’s performance in modulating his brain state in relation, not only to the target stimulus but also the other base stimuli whose dissimilarities generate the RS. In practice, the fewer the steps (i.e. the number of task blocks) to reach the target, the faster the dissimilarity with the target brain state decreases and the dissimilarities with the other base stimuli replicate the original ones. Furthermore, estimating the distance between the projection of the last block and the target provides a measure of success for the NF outcome, as the lower this distance, the better the participant has likely fulfilled the task of engaging in that specific mental representation.

#### 2.8.3 Simulations

An ensemble of artificial datasets was created to simulate the outcome of the rt-RSA analysis under different noise conditions.

For each simulation, five different artificial time-series of 430 images (the same number of time points of the real localizer data) were simulated in an ROI of 100 voxels. All voxel timeseries of all data sets were initialized with random Gaussian noise with zero mean and different variances (σ = 0.5, 1, 2, 3). In each data set, an ideal activation time-course was injected into a variable percentage of voxels, whereas the rest of the voxels only contained noise. To simulate five different patterns (associated with five simulated stimuli), the subsets of active voxels were varied across the five artificial data sets. In particular, assigning a numerical index to the five patterns from 0 to 4, the percentage of active voxels for the even patterns was randomly extracted from a uniform distribution of integers ranging from 30 to 60, whereas the percentage of active voxels for the odd patterns was randomly extracted from a uniform distribution of integers ranging from 20 to 50. The activation time course was generated by convolving the canonical hemodynamic response function with a box-car function according to the same paradigm of the real localizer experiment, however, the signal amplitude of each block was scaled with a random factor from a uniform distribution ranging from −1 to 3, to simulate variable modulation performances of the participant across task blocks.

The procedure described in the previous paragraph was repeated 1000 times, and for each simulation, the distances of the projected brain activity in the RS from the target, as well as the correlation of the RDM calculated at the end of each task block with the reference RDM, were calculated. These performances were thus reported on the average of 1000 simulations. The whole analysis was repeated four times after increasing level of noise variance (σ = 0.5, 1, 2, 3) to evaluate the performance of the rt-RSA under various noise conditions.

### 3. RESULTS

The localizer data were preliminary used to simulate a different NF experiment for each stimulus, by extracting the brain pattern of the ROI at the end of each task block and calculating its position in the RS. The estimated distance between each projection (as obtained via incremental online GLM) and its corresponding target (as obtained via full offline GLM) showed, for all the stimuli, a decreasing trend over task blocks and a value of zero at the end of the session. In particular, the stimulus “cat” showed a maximum distance from the target of 0.6 at the first task block and a distance lower than 0.2 from the 6^th^ task block on. The stimulus “dog” showed a maximum distance from the target of 0.5 at the first task block and a distance lower than 0.2 from the 8^th^ task-block on. Similarly, the stimulus “hammer” showed a maximum distance from the target of 0.65 at the 2^nd^ task block and a distance lower than 0.2 from the 8^th^ task-block on. The stimulus “chair” showed more variability with a maximum distance of 1.2 at the 3^rd^ task-block and a distance lower than 0.2 only at the 10^th^ task block (Figure 3).

**Figure 3.**
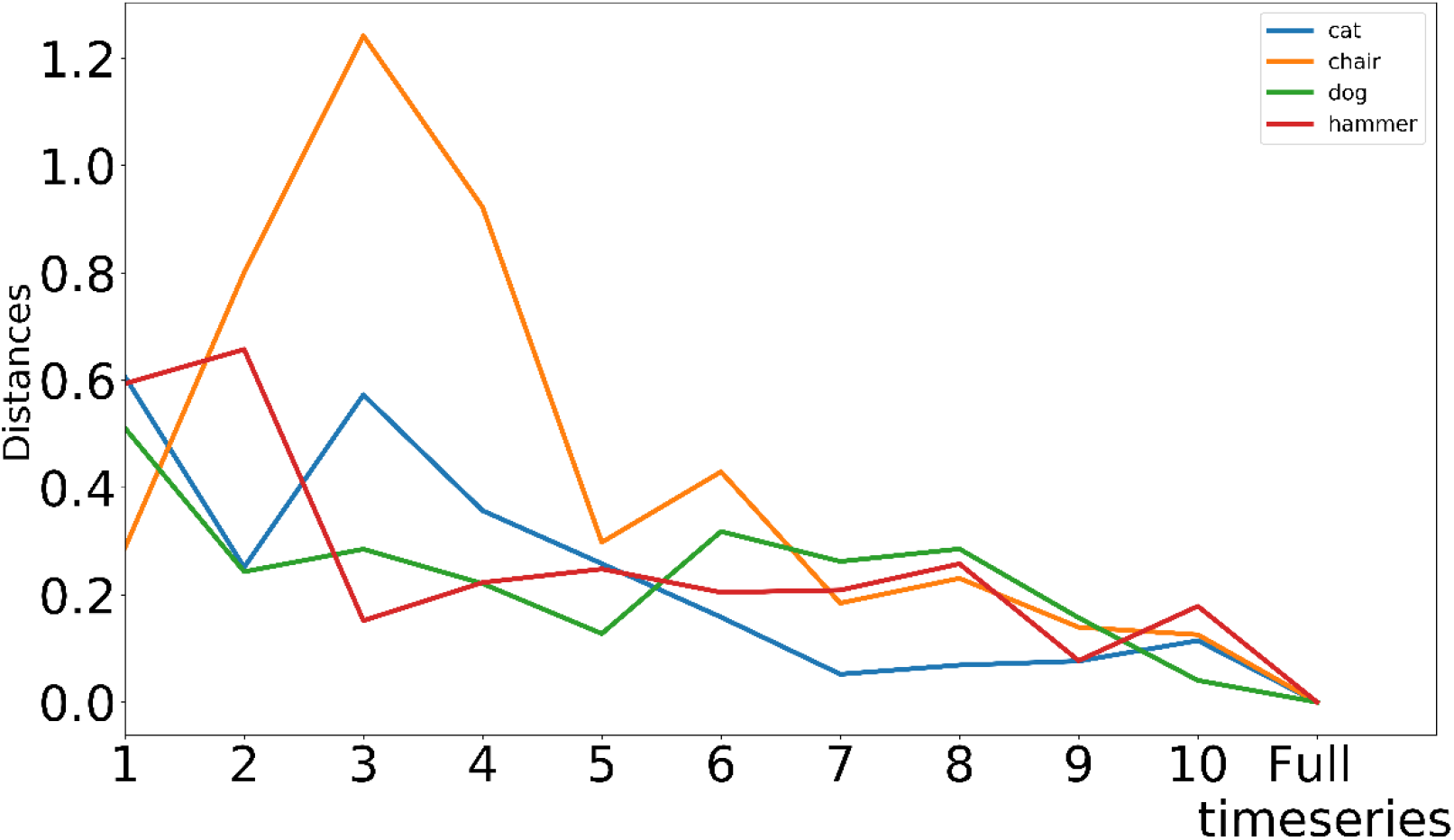
Distance of the projections from the target in the RS over the task-blocks, evaluated on the localizer data. Four neurofeedback experiments were simulated sampling incrementally the fMRI data of one of the stimuli and projecting these values in the defined RS. The estimated distances have a decreasing trend for all the stimuli and a zero value in the last task-block.

The trajectories of the projections in the RS showed, for all the stimuli, that, at the end of the time series, the positions of the estimated brain patterns coincide with the positions of the corresponding targets. In addition, the visual inspection of the trajectories suggested how each stimulus follows a different path towards its target (Figure 4).

**Figure 4.**
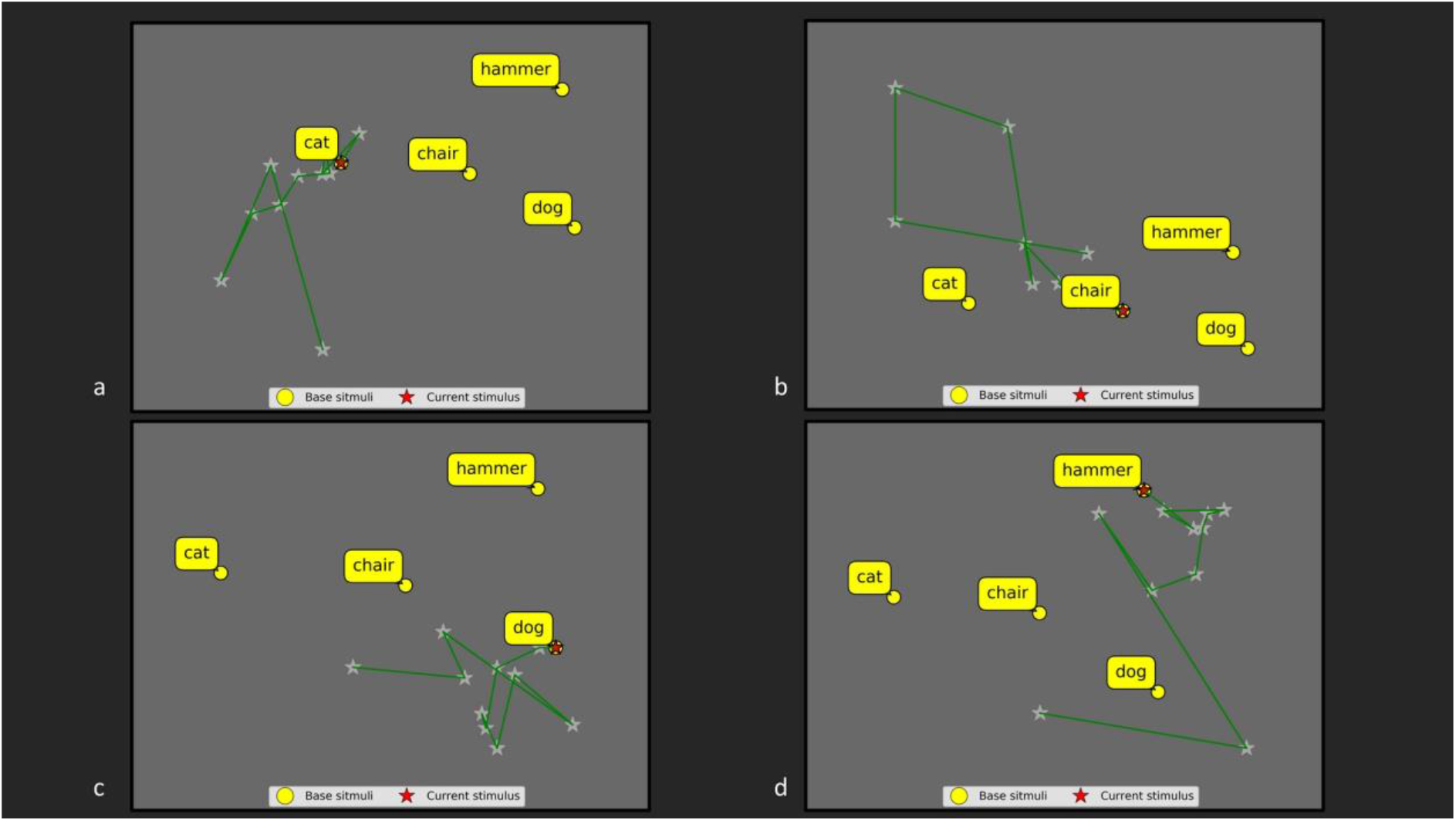
Trajectories of the projections on the RS over the task-blocks, evaluated on the localizer data. Four neurofeedback experiments were simulated by sampling incrementally the fMRI data of one of the stimuli and projecting these values in the defined RS. The estimated projections for all the stimuli move towards (and at the end perfectly overlap with) the corresponding targets.

The performances of the subject during the real NF experiment were evaluated using the data from both the localizer and the NF session.

The Spearman correlation between RDMs, estimated at each task block, and the reference RDM, estimated at the end of the localizer session with a full GLM, showed an overall increasing trend across the task blocks (Figure 5a). The correlation coefficient remained negative from the 1^st^ to the 4^th^ task block, before turning positive at the 5 ^th^ task block, reaching a maximum of 0.8 at the 7^th^ task block, and, finally, falling down to lower values (~0.4) in the last two task blocks (Figure 5a). The estimated distances from the target in the RS showed an overall decreasing trend across the task-blocks. In the first four task blocks, the distances remained higher than 0.4 with a maximum of 0.82 at the 3^rd^ task block, but then the distance from the target decreased towards a minimum of 0.12 at the last time-point (Figure 5b).

**Figure 5.**
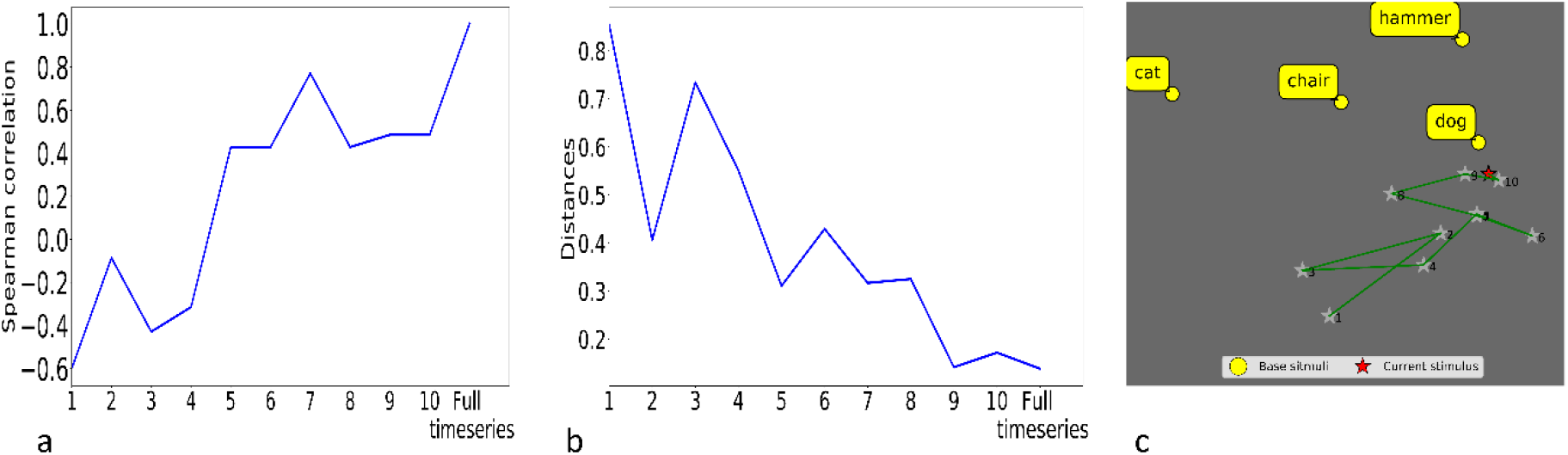
Evaluation of the subject’s performance: correlation between RDMs in the localizer data (a), distances between the projection and the target in the RS (b) and trajectory of the projection in the RS (c). The Spearman’s correlation between the RDMs estimated at the end of each task-block (i.e. a subset of the time series) and the reference RDM estimated at the end of the time series, show an increasing trend suggesting a stabilization of the brain patterns in the localizer data (a). The estimated distances between the projection in the RS and the target show a decreasing trend supporting the idea that, in the neurofeedback session, the subject learned to modulate his brain pattern in order to engage in the target mental state (b). The trajectories of the projections in the RS confirm that the subject managed to move the point corresponding to his brain activity towards the target (i.e. “dog”) (c).

The analyses of the artificial datasets were useful to evaluate the stability and accuracy of the rt-RSA under different noise conditions by estimating the distances from the projection of the current brain activity (at each task block) to the target in the RS and the correlation between the RDMs at each task block and the reference RDM. After 1000 simulations, the average distance exhibited a monotonic decreasing trend over time for different noise conditions (Figure 6) with a minimum of ~0.05 for *σ* = 0.5 and a maximum of ~0.2 for *σ* = 3. An opposite trend was observed in the analysis of the RDM correlations. Namely, the average Spearman correlation coefficient exhibited a monotonic increasing trend over time for each noise level. The exponential nature of the curve was more evident at lower noise levels (σ = 0.5, 1). At the first block, the starting point of the curve showed lower values at higher noise levels, ranging from a maximum of 0.825 (σ = 0.5) to a minimum of about 0.2 (σ = 3).

**Figure 6.**
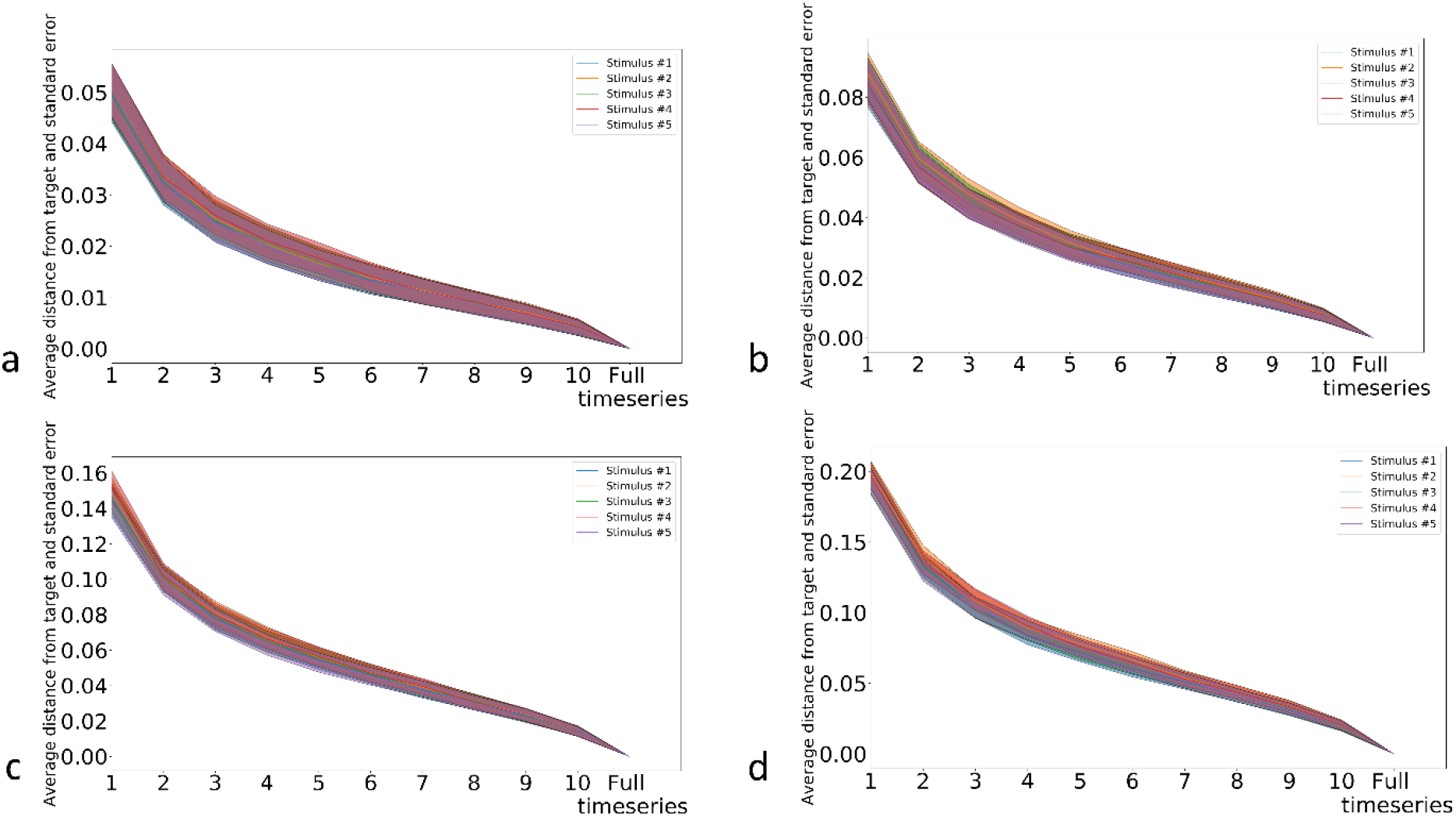
Distances from the target in the RS on the average on 1000 simulated datasets in various noise conditions: data with gaussian noise μ =0, *σ* = 0.5 (a), data with gaussian noise μ = 0, *σ* = 1 (b),data with gaussian noise μ =0, *σ* = 2 (c) and data with gaussian noise μ = 0, *σ* = 3 (d). In all the cases, the distances from the target have a decreasing trend towards zero.

## 4. DISCUSSION

In this work, rt-RSA, a method based on RSA and a novel procedure for the real-time embedding of the current mental state in the 2D visualization via cMDS, has been introduced for rt-fMRI-NF paradigms. The online implementation of RSA (in a real-time incremental procedure) enabled a novel type of multi-dimensional feedback of the participant’s brain activity which can be semantically related to an internal stimulus representation (thereby reflecting the actual mental state of the subject) via multi-voxel pattern analysis. Using rt-RSA, the online estimated neural pattern is displayed as a movable point on a plane and engagement in mental tasks navigates the point towards (or away from) two or more target points displayed at fixed positions. The relative distance between the current location of the NF point – representing the current mental state – and the other points approximate their mutual (dis)similarities. The other semantic “anchor” points reflect mental states that the participant was previously engaged in during a localizer session. In this way, the participant is requested to change the position of the movable point towards one of the target points in a fixed RS space by self-modulating his/her own brain activity until reaching a desired mental state.

There are (at least) two potential advantages of this framework over classical NF paradigms: first, the NF display encodes (and therefore the self-modulation operates on) multi-voxel (i.e. spatially distributed) patterns of brain activity within one or more selected ROIs, which are (known or found to be) critically involved in the semantic processing of the stimulus, rather than one-dimensional (spatially averaged) ROI signals [17,23,24] or pairwise connectivity estimates [25]; second, this (locally) distributed pattern of brain activity is dynamically compared, not just to one, but many, base or reference patterns, thus providing additional degrees of freedom, both to the experimenter (in the preparation and set-up of the NF materials) and to the participant (in the choice of the mental strategy), to more efficiently (self-) modulate a mental representation along multiple dimensions.

Rt-RSA was successfully tested in a real rt-fMRI-NF experiment at 7 Tesla where the subject performed a visual imagery task and the brain activity from an ROI in the left ITC was recorded.

The imagery task was chosen for a proof-of-concept study, as it enabled the possibility to modulate the brain activity related to semantic features of the stimuli in a region known to encode the high-level representation of natural objects [42,55,56]. The functional data from a localizer session were used to both generate the RS for a set of four base stimuli (to be used in the NF session) and to simulate NF experiments. While the actual NF performances of the subject were evaluated on the real NF data from the NF session, the computational feasibility and the statistical accuracy of the rt-RSA approach under simulated ideal conditions of successful modulation were evaluated on the same localizer data. In addition, 1000 simulations of multi-voxel ROI patterns were performed to evaluate the rt-RSA approach under different signal-to-noise conditions.

The analysis of the simulated NF experiments with real fMRI data from the localizer session demonstrated that it is possible to implement a visual two-dimensional NF by estimating in real-time the current brain pattern by dynamically updating (with negligible delays) the position of the associated current mental representation in a previously defined (and fixed) RS. Indeed, the resulting trajectory of a point on a two-dimensional plane (which is obtained by analysing incrementally the fMRI time-series for just one target stimulus at the time) and the final collapsing of this point to the target position (when the full time-series is used in the GLM) demonstrated that the provided visual feedback could be in principle correct and stable. More specifically, the results of the simulated NF experiments clearly showed that if the participant is in principle able to become engaged in the same mental state which causes exactly the same distribution of brain activity elicited by the target stimulus in the target ROI, the resulting visual feedback could in principle guide this process towards the perfect match between the positions of the current and target brain patterns.

The results from the rt-fMRI-NF experiment demonstrate that the rt-RSA approach can be applied in a real-world scenario, with little additional computational steps and no additional hardware requirements. Thus, our proof-of-concept study suggests that it would be possible to integrate an rt-RSA based procedure within several experiments using existing NF paradigms. The integration of a multi-dimensional NF display did not introduce additional difficulties for the participant to understand the task, at least according to the report from the single participant examined in this study. Indeed, while the task progress is perceived by the subject like “a journey in a geographical map”, the investigator can still rely on the distance from the target as a simple (one-dimensional) index of success to evaluate the subject’s performance [19]. Moreover, the offline analyses performed on the localizer data showed how it is possible to obtain a preliminary assessment of the stability of the RS as initially created from the base stimuli, in a similar way as it happens in a one-dimensional NF where it is possible to evaluate the variability of the BOLD percent signal changes over the trials.

The analyses of the artificial datasets allowed to assess the impact of noise on the trajectories under ideal conditions of successful NF. These gathered two main observations: First, the projection of the current neural representation, and its final convergence to the target points, remains stable also when the fMRI signal is affected by a relatively higher amount of noise. However, higher noise levels critically affect the position of the current point at the beginning of the experiment, thereby potentially increasing the length of the trajectory in the RS, and therefore the minimum number of steps required, to reach the target. Second, the amount of noise has an impact on the reference RDM itself and, therefore, on the RS, thereby the monotonic correlations between current and reference RDMs may be reduced.

Taken together, these results may suggest that, in an ideal scenario, the presence of higher noise levels simply causes longer paths to the target state, i.e. noise itself does not necessarily undermine or disrupt the stability of the rt-RSA under the assumption that the participant has successfully learnt to regulate his/her mental states with the decided strategy. However, it should be also pointed out that the stabilization of the statistical values does not represent per se an indication that the subject has successfully consolidated an optimal mental strategy. Actually, if the subject chooses a wrong strategy, there is no guarantee of successful convergence, even at very low noise levels. Nonetheless, as far as the noise is assumed constant across the blocks, the final distance of the current, from the target, representation may still provide a useful indication of the overall performance in modulating the mental state according to that strategy.

The proposed method may become a promising tool for the self-regulation of brain signals as it is flexible and versatile and provides both the participant and experimenter with transparent visual feedback about how the current mental state relates to related “anchor” mental states in a semantic map. There are, however, no fundamental limitations about the feedback modality as different physical dimensions can be used as different feedback channels. The choice of the mental task, for which no detailed instructions are needed, is completely up to the participants and their own representations, the only requirement being that a robust activation pattern is elicited (and verified) during the localizer session. As a consequence, besides integrating a visual feedback during object imagery, as shown in the present study, rt-RSA could be in principle used in more complex scenarios such as those employed in NF-based emotion regulation [11]. In fact, basic (positive or negative) emotions have been previously associated with specific neural signatures within different brain regions [59], including, e.g., the amygdala [60]. Therefore, it could be possible to project (and target) patterns with different emotional valence, on the same RS from one (or more) of these brain regions, after training the subject to engage in some different mental states that become associated with different local patterns [11] during the localizer session. Technically, it is only essential to keep the same number and order of voxels in the ROI(s) to correctly fit the original dimensions of the RS [32].

Finally, towards a proper generalization of this approach, and to counteract the inter-subject variability in local patterns, as resulting from the localizing/training phase with more complex stimuli, it could be possible to force higher levels of smoothing to the functional data and, in principle, derive one unique RDM from the group-level analysis of several subjects (e.g. from a healthy or control cohort). Along these lines, it would be possible to create one “external” RS to be used in the NF session of a single individual (e.g. a patient) by aligning the estimated neural pattern from the individual to a common space [61,62]. As a consequence, the participants could be asked to self-modulate their own mental state by navigating through predefined mental states associated, not (only) with their own neural representations, but rather with some “control” representations from different people. However, this approach will need proper design and careful testing to be validated prior to any clinical applications.

## 5. CONCLUSION

In conclusion, a new method for rt-fMRI-NF has been introduced which promises to go beyond the classical approach of fMRI signal self-modulation. The presented simulation and the preliminary results from an rt-fMRI-NF proof-of-concept study demonstrate that rt-RSA provides the possibility to project the current mental state in a fixed representational space for a given ROI enabling a semantic feedback to the subject for the self-regulation of mental states in a multi-dimensional space. This approach has been shown to be computationally and statistically feasible and its application to a real NF experiment based on a simple imagery task at 7 Tesla has yielded encouraging results. Nonetheless, future studies are warranted to increase the number of subjects and to assess performances in a multi-subject study, eventually at lower (i.e. clinical) magnetic fields (e.g. 1.5 or 3 Tesla).

**Figure 7.**
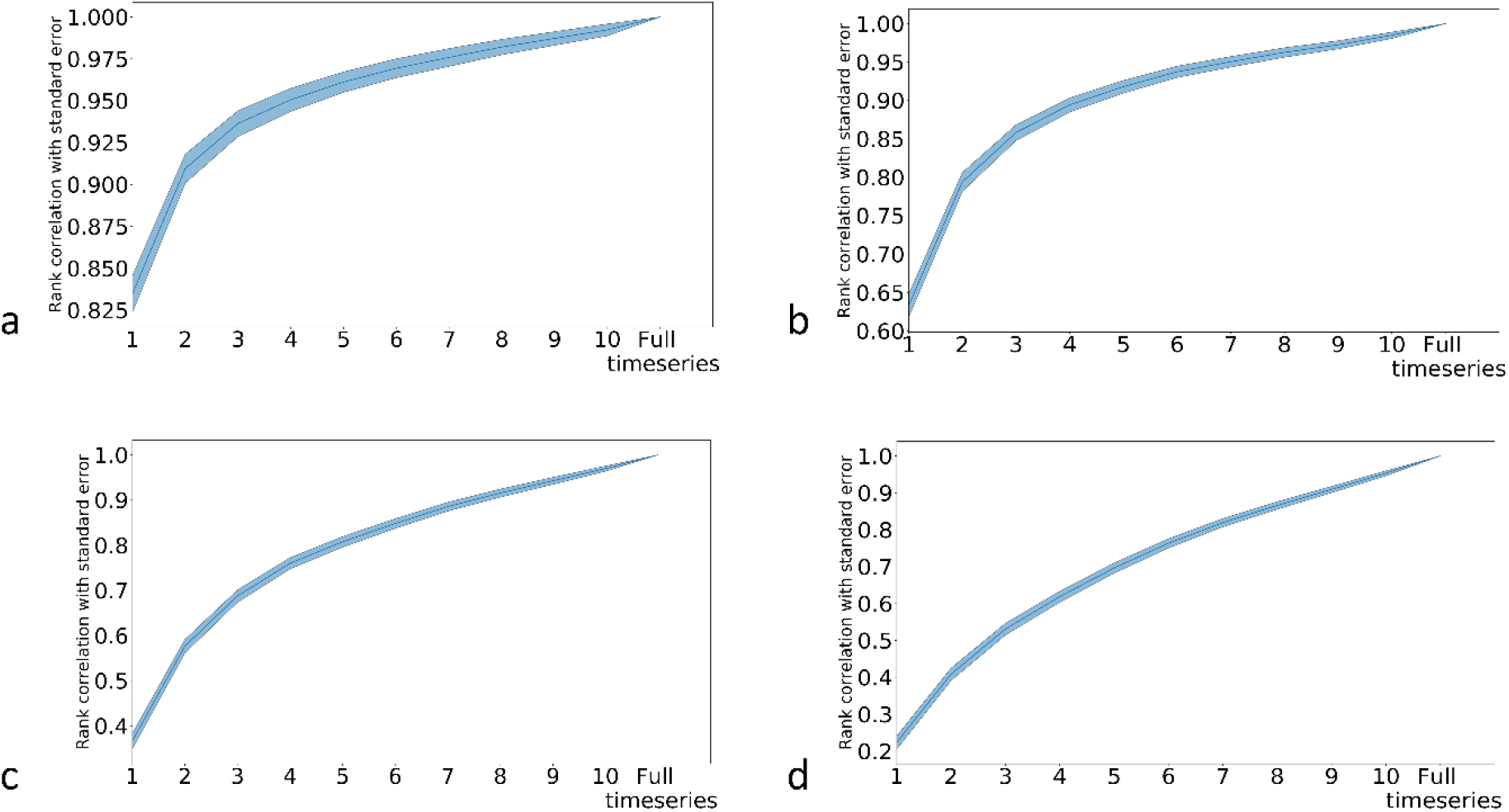
Spearman’s correlation between RDMs estimated at the end of each task-block and the reference RDM on the average on 1000 simulated datasets in various noise conditions: data with gaussian noise μ =0, *σ*= 0.5 (a), data with gaussian noise μ =0, *σ*= 1 (b),data with gaussian noise μ =0, *σ*= 2 (c) and data with gaussian noise μ =0, *σ*= 3 (d). In all the cases, the correlations with the reference RDM have a increasing exponential trend towards one.

## Acknowledgments

Icons for the Figure 1 and Figure 2: “brain activity” by Ben Davis, “Computer” by iconcheese, “dots” by Alexander Skowalsky, “BCG matrix” by Kirby Wu, “scatter chart” by KonKapp, “Sound” by Adrien Coquet and “MRI scanner” by Healthcare Symbols were distributed by https://thenounproject.com/ under Creative Commons CCBY license.

## References

[1] Cohen Kadosh K, Luo Q, de Burca C, Sokunbi M O, Feng J, Linden D E J and Lau J Y F 2016 Using real-time fMRI to influence effective connectivity in the developing emotion regulation network Neuroimage 125 616–26

[2] Yamin H G, Gazit T, Tchemodanov N, Raz G, Jackont G, Charles F, Fried I, Hendler T and Cavazza M 2017 Depth electrode neurofeedback with a virtual reality interface Brain-Computer Interfaces 4 201–13

[3] Ramot M, Grossman S, Friedman D and Malach R 2016 Covert neurofeedback without awareness shapes cortical network spontaneous connectivity PNAS 113 E2413–20

[4] Buch E, Weber C, Cohen L G, Braun C, Dimyan M A, Ard T, Mellinger J, Caria A, Soekadar S, Fourkas A and Birbaumer N 2008 Think to Move: a Neuromagnetic Brain-Computer Interface (BCI) System for Chronic Stroke Stroke 39 910–7

[5] Young B M, Nigogosyan Z, Walton L M, Remsik A, Song J, Nair V A, Tyler M E, Edwards D F, Caldera K, Sattin J A, Williams J C and Prabhakaran V 2015 Dose-response relationships using brain–computer interface technology impact stroke rehabilitation Front Hum Neurosci 9

[6] Sitaram R, Ros T, Stoeckel L, Haller S, Scharnowski F, Lewis-Peacock J, Weiskopf N, Blefari M L, Rana M, Oblak E, Birbaumer N and Sulzer J 2017 Closed-loop brain training: the science of neurofeedback Nature Reviews Neuroscience 18 86–100

[7] Watanabe T, Sasaki Y, Shibata K and Kawato M 2017 Advances in fMRI Real-Time Neurofeedback Trends in Cognitive Sciences 21 997–1010

[8] Scharnowski F and Weiskopf N 2015 Cognitive enhancement through real-time fMRI neurofeedback Current Opinion in Behavioral Sciences 4 122–7

[9] Sitaram R, Veit R, Stevens B, Caria A, Gerloff C, Birbaumer N and Hummel F 2011 Acquired Control of Ventral Premotor Cortex Activity by Feedback Training: An Exploratory Real-Time fMRI and TMS Study Neurorehabilitation and Neural Repair

[10] Herwig U, Lutz J, Scherpiet S, Scheerer H, Kohlberg J, Opialla S, Preuss A, Steiger V R, Sulzer J, Weidt S, Stämpfli P, Rufer M, Seifritz E, Jäncke L and Brühl A B 2019 Training emotion regulation through real-time fMRI neurofeedback of amygdala activity NeuroImage 184 687–96

[11] Linhartová P, Látalová A, Kóša B, Kašpárek T, Schmahl C and Paret C 2019 fMRI neurofeedback in emotion regulation: A literature review NeuroImage 193 75–92

[12] Rota G, Sitaram R, Veit R, Erb M, Weiskopf N, Dogil G and Birbaumer N 2009 Self-regulation of regional cortical activity using real-time fmri: the right inferior frontal gyrus and linguistic processing Human Brain Mapping 30 1605–14

[13] Shibata K, Watanabe T, Sasaki Y and Kawato M 2011 Perceptual Learning Incepted by Decoded fMRI Neurofeedback Without Stimulus Presentation Science 334 1413–5

[14] Linden D E J, Habes I, Johnston S J, Linden S, Tatineni R, Subramanian L, Sorger B, Healy D and Goebel R 2012 Real-Time Self-Regulation of Emotion Networks in Patients with Depression PLoS One 7

[15] Mehler D M A, Sokunbi M O, Habes I, Barawi K, Subramanian L, Range M, Evans J, Hood K, Lührs M, Keedwell P, Goebel R and Linden D E J 2018 Targeting the affective brain—a randomized controlled trial of real-time fMRI neurofeedback in patients with depression Neuropsychopharmacology 43 2578–85

[16] Orlov N D, Giampietro V, O’Daly O, Lam S-L, Barker G J, Rubia K, McGuire P, Shergill S S and Allen P 2018 Real-time fMRI neurofeedback to down-regulate superior temporal gyrus activity in patients with schizophrenia and auditory hallucinations: a proof-of-concept study Translational Psychiatry 8 1–10

[17] Young K D, Zotev V, Phillips R, Misaki M, Yuan H, Drevets W C and Bodurka J 2014 Real-Time fMRI Neurofeedback Training of Amygdala Activity in Patients with Major Depressive Disorder PLoS One 9

[18] Zweerings J, Hummel B, Keller M, Zvyagintsev M, Schneider F, Klasen M and Mathiak K 2019 Neurofeedback of core language network nodes modulates connectivity with the default-mode network: A double-blind fMRI neurofeedback study on auditory verbal hallucinations NeuroImage 189 533–42

[19] Paret C, Goldway N, Zich C, Keynan J N, Hendler T, Linden D and Cohen Kadosh K 2019 Current progress in real-time functional magnetic resonance-based neurofeedback: Methodological challenges and achievements NeuroImage 202 116107

[20] Kober S E, Witte M, Ninaus M, Neuper C and Wood G 2013 Learning to modulate one’s own brain activity: the effect of spontaneous mental strategies Front Hum Neurosci 7

[21] Zilverstand A, Sorger B, Sarkheil P and Goebel R 2015 fMRI neurofeedback facilitates anxiety regulation in females with spider phobia Front Behav Neurosci 9 148

[22] Amano K, Shibata K, Kawato M, Sasaki Y and Watanabe T 2016 Learning to associate orientation with color in early visual areas by associative decoded fMRI neurofeedback Curr Biol 26 1861–6

[23] deCharms R C, Christoff K, Glover G H, Pauly J M, Whitfield S and Gabrieli J D E 2004 Learned regulation of spatially localized brain activation using real-time fMRI NeuroImage 21 436–43

[24] Weiskopf N, Veit R, Erb M, Mathiak K, Grodd W, Goebel R and Birbaumer N 2003 Physiological self-regulation of regional brain activity using real-time functional magnetic resonance imaging (fMRI): Methodology and exemplary data NeuroImage 19 577–86

[25] Koush Y, Rosa M J, Robineau F, Heinen K, W. Rieger S, Weiskopf N, Vuilleumier P, Van De Ville D and Scharnowski F 2013 Connectivity-based neurofeedback: Dynamic causal modeling for real-time fMRI Neuroimage 81 422–30

[26] Cao M, Wang J-H, Dai Z-J, Cao X-Y, Jiang L-L, Fan F-M, Song X-W, Xia M-R, Shu N, Dong Q, Milham M P, Castellanos F X, Zuo X-N and He Y 2014 Topological organization of the human brain functional connectome across the lifespan Developmental Cognitive Neuroscience 7 76–93

[27] Fair D A, Dosenbach N U F, Church J A, Cohen A L, Brahmbhatt S, Miezin F M, Barch D M, Raichle M E, Petersen S E and Schlaggar B L 2007 Development of distinct control networks through segregation and integration Proc Natl Acad Sci U S A 104 13507–12

[28] Shibata K, Lisi G, Cortese A, Watanabe T, Sasaki Y and Kawato M 2019 Toward a comprehensive understanding of the neural mechanisms of decoded neurofeedback NeuroImage 188 539–56

[29] Krause F, Benjamins C, Lührs M, Eck J, Noirhomme Q, Rosenke M, Brunheim S, Sorger B and Goebel R 2017 Real-time fMRI-based self-regulation of brain activation across different visual feedback presentations Brain-Computer Interfaces 4 87–101

[30] Kriegeskorte N, Mur M and Bandettini P 2008 Representational similarity analysis - connecting the branches of systems neuroscience Frontiers in Systems Neuroscience 2

[31] Dennett D C 1987 The Intentional Stance (The MIT Press)

[32] Kriegeskorte N and Kievit R A 2013 Representational geometry: Integrating cognition, computation, and the brain Trends in Cognitive Sciences 17 401–12

[33] Nili H, Wingfield C, Walther A, Su L, Marslen-Wilson W and Kriegeskorte N 2014 A Toolbox for Representational Similarity Analysis PLoS Computational Biology 10

[34] Wilcoxon F 1945 Individual Comparisons by Ranking Methods Biometrics Bulletin 1 80–3

[35] Walther A, Nili H, Ejaz N, Alink A, Kriegeskorte N and Diedrichsen J 2016 Reliability of dissimilarity measures for multi-voxel pattern analysis NeuroImage 137 188–200

[36] Borg I and Groenen P J F 2005 Modern Multidimensional Scaling: Theory and Applications (New York: Springer-Verlag)

[37] Kruskal J B and Wish M 1978 Multidimensional Scaling (SAGE)

[38] Wang J 2012 Classical Multidimensional Scaling Geometric Structure of High-Dimensional Data and Dimensionality Reduction ed J Wang (Berlin, Heidelberg: Springer) pp 115–29

[39] Brouwer G J and Heeger D J 2009 Decoding and Reconstructing Color from Responses in Human Visual Cortex J. Neurosci. 29 13992–4003

[40] Op de Beeck H P, Torfs K and Wagemans J 2008 Perceived Shape Similarity among Unfamiliar Objects and the Organization of the Human Object Vision Pathway Journal of Neuroscience 28 10111–23

[41] Giordano B L, McAdams S, Zatorre R J, Kriegeskorte N and Belin P 2013 Abstract Encoding of Auditory Objects in Cortical Activity Patterns Cereb Cortex 23 2025–37

[42] Kriegeskorte N, Mur M, Ruff D A, Kiani R, Bodurka J, Esteky H, Tanaka K and Bandettini P A 2008 Matching Categorical Object Representations in Inferior Temporal Cortex of Man and Monkey Neuron 60 1126–41

[43] Naselaris T, Stansbury D E and Gallant J L 2012 Cortical representation of animate and inanimate objects in complex natural scenes Journal of Physiology-Paris 106 239–49

[44] Polyn S M, Natu V S, Cohen J D and Norman K A 2005 Category-Specific Cortical Activity Precedes Retrieval During Memory Search Science 310 1963–6

[45] Wiestler T, McGonigle D J and Diedrichsen J 2011 Integration of sensory and motor representations of single fingers in the human cerebellum Journal of Neurophysiology 105 3042–53

[46] Torgerson W S 1958 Theory and methods of scaling (Wiley)

[47] de Silva V and Tenenbaum J B Sparse multidimensional scaling using landmark points 41

[48] Krause F and Lindemann O 2014 Expyriment: A Python library for cognitive and neuroscientific experiments Behav Res 46 416–28

[49] Hebart M N, Dickter A H, Kidder A, Kwok W Y, Corriveau A, Van Wicklin C and Baker C I 2019 THINGS: A database of 1,854 object concepts and more than 26,000 naturalistic object images ed F A Soto PLOS ONE 14 e0223792

[50] Peirce J, Gray J R, Simpson S, MacAskill M, Höchenberger R, Sogo H, Kastman E and Lindeløv J K 2019 PsychoPy2: Experiments in behavior made easy Behav Res 51 195–203

[51] Feinberg D A, Moeller S, Smith S M, Auerbach E, Ramanna S, Glasser M F, Miller K L, Ugurbil K and Yacoub E 2010 Multiplexed echo planar imaging for sub-second whole brain fmri and fast diffusion imaging PLoS ONE 5

[52] Moeller S, Yacoub E, Olman C A, Auerbach E, Strupp J, Harel N and Uğurbil K 2010 Multiband Multislice GE-EPI at 7 Tesla, With 16-Fold Acceleration Using Partial Parallel Imaging With Application to High Spatial and Temporal Whole-Brain FMRI Magnetic Resonance in Medicine 63 1144–53

[53] Xu J, Moeller S, Auerbach E J, Strupp J, Smith S M, Feinberg D A, Yacoub E and Uğurbil K 2013 Evaluation of slice accelerations using multiband echo planar imaging at 3 Tesla NeuroImage 83 790–5

[54] Boynton G M, Engel S A, Glover G H and Heeger D J 1996 Linear Systems Analysis of Functional Magnetic Resonance Imaging in Human V1 J Neurosci 16 4207–21

[55] Haxby J V, Gobbini M I, Furey M L, Ishai A, Schouten J L and Pietrini P 2001 Distributed and Overlapping Representations of Faces and Objects in Ventral Temporal Cortex Science 293 2425–30

[56] Mur M, Meys M, Bodurka J, Goebel R, Bandettini P A and Kriegeskorte N 2013 Human Object-Similarity Judgments Reflect and Transcend the Primate-IT Object Representation Front. Psychol. 4

[57] Yarkoni T, Poldrack R A, Nichols T E, Van Essen D C and Wager T D 2011 Large-scale automated synthesis of human functional neuroimaging data Nat Methods 8 665–70

[58] Genovese C R, Lazar N A and Nichols T 2002 Thresholding of statistical maps in functional neuroimaging using the false discovery rate NeuroImage 15 870–8

[59] Saarimäki H, Gotsopoulos A, Jääskeläinen I P, Lampinen J, Vuilleumier P, Hari R, Sams M and Nummenmaa L 2016 Discrete Neural Signatures of Basic Emotions Cereb Cortex 26 2563–73

[60] Sergerie K, Chochol C and Armony J L 2008 The role of the amygdala in emotional processing: A quantitative meta-analysis of functional neuroimaging studies Neuroscience & Biobehavioral Reviews 32 811–30

[61] Frost M A and Goebel R 2013 Functionally informed cortex based alignment: An integrated approach for whole-cortex macro-anatomical and ROI-based functional alignment NeuroImage 83 1002–10

[62] Haxby J V, Guntupalli J S, Connolly A C, Halchenko Y O, Conroy B R, Gobbini M I, Hanke M and Ramadge P J 2011 A common, high-dimensional model of the representational space in human ventral temporal cortex Neuron 72 404–16

